# Modulation of RNA binding properties of the RNA helicase UPF1 by its activator UPF2

**DOI:** 10.1101/2022.03.26.485946

**Authors:** Guangpu Xue, Vincent D. Maciej, Alexandrina Machado de Amorim, Melis Pak, Uma Jayachandran, Sutapa Chakrabarti

**Author notes:** To whom correspondence should be addressed: Sutapa Chakrabarti.

## Abstract

The NMD helicase UPF1 is a prototype of the superfamily 1 (SF1) of RNA helicases that bind RNA with high affinity and translocate on it in an ATP-dependent manner. Previous studies showed that UPF1 has a low basal catalytic activity that is greatly enhanced upon binding of its interaction partner, UPF2. Activation of UPF1 by UPF2 entails a large conformational change that switches the helicase from an RNA-clamping mode to an RNA-unwinding mode. The ability of UPF1 to bind RNA was expected to be unaffected by this activation mechanism. Here we show, using a combination of biochemical and biophysical methods, that binding of UPF2 to UPF1 drastically reduces the affinity of UPF1 for RNA, leading to a release of the bound RNA. Although UPF2 is capable of binding RNA *in vitro*, our results suggest that dissociation of the UPF1-RNA complex is not a consequence of direct competition in RNA binding but rather an allosteric effect that is likely mediated by the conformational change induced by binding of the helicase to its activator. We discuss these results in light of transient interactions forged during mRNP assembly, particularly in the UPF1-dependent mRNA decay pathways.

## Introduction

The coat of proteins that assemble on a messenger RNA (mRNA), leading to the formation of a messenger ribonucleoprotein (mRNP) represent a major class of effectors of post-transcriptional gene regulation (Gehring, Wahle et al. 2017). Messenger RNPs are often transient in nature, and undergo rapid assembly and disassembly depending on the functional requirement of the cell. A class of proteins that mediate remodeling while being an integral part of mRNPs themselves are RNA helicases. These enzymes are molecular motors that harness the energy of ATP binding and hydrolysis to bring about conformational changes in RNA, which facilitate binding of some protein factors while preventing others from associating with the RNA. RNA helicases can also mediate protein-protein interactions, often acting as a scaffold for mRNP assembly (Linder and Jankowsky 2011, Bourgeois, Mortreux et al. 2016).

The RNA helicase Up-Frameshift1 (UPF1) is a central effector of the nonsense-mediated mRNA decay (NMD) pathway and has diverse functions, ranging from target selection in the early steps of the pathway to mRNP remodeling and recruitment of additional protein factors in the later stages of NMD (Leeds, Peltz et al. 1991, Ohnishi, Yamashita et al. 2003, Okada-Katsuhata, Yamashita et al. 2012, Lee, Pratt et al. 2015). The catalytic activity of UPF1 is greatly enhanced upon binding to a core NMD factor UPF2, which also serves as an adaptor that connects the helicase to the exon-junction complex (EJC) (Chamieh, Ballut et al. 2008, Buchwald, Ebert et al. 2010). UPF1 is a multi-domain protein, consisting of a cis-inhibitory cysteine-histidine rich (CH) domain and a helicase core comprising two RecA-like domains and two auxiliary domains, 1B and 1C (Figure 1A) (Cheng, Muhlrad et al. 2007, Chamieh, Ballut et al. 2008). The RecA-like domains together make up a deep cleft for ATP binding and on the opposite side, a shallow surface which allows RNA to bind across it. UPF1 spans 9-11 nucleotides on single-stranded RNA and binds with a high affinity of approximately 50 nM (Chakrabarti, Jayachandran et al. 2011). The basal catalytic activity of UPF1 is repressed owing to clamping down of the auxiliary domain 1B on the 3’-end of the RNA. The clamp is positioned indirectly by the CH domain, which interacts with the RecA2 domain to hold the helicase in an inactive conformation. Activation of UPF1 is due to a large conformational change in the helicase that is brought about upon binding of UPF2 to the CH domain (Clerici, Mourao et al. 2009, Chakrabarti, Jayachandran et al. 2011).

**Figure 1.**
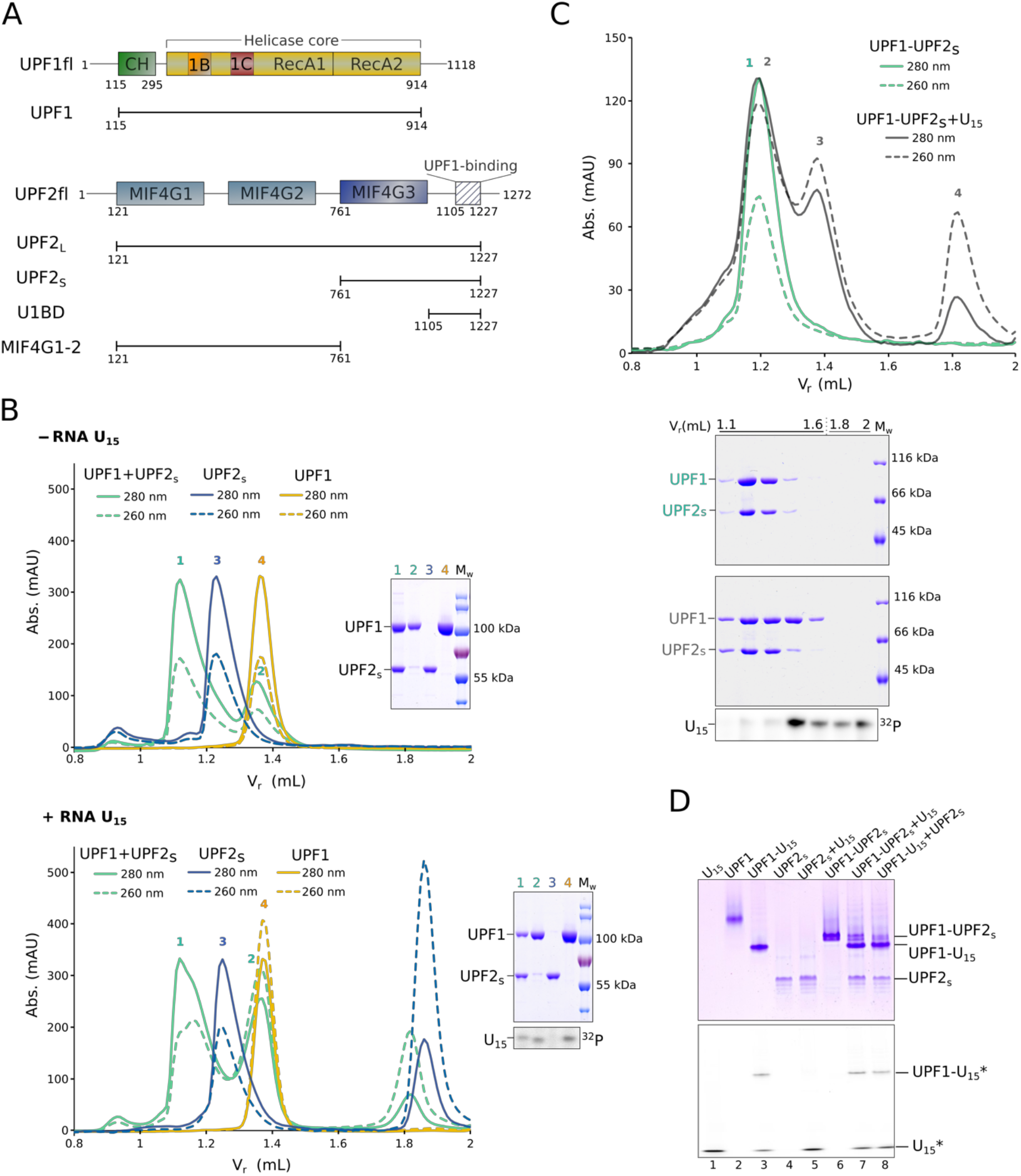
UPF1 does not engage in concomitant interactions with UPF2 and RNA. A) Schematic of the domain organization of UPF1 and UPF2. The helicase core comprising the RecA1 and RecA2 domains is colored yellow. The CH domain and auxiliary domains 1B and 1C are shown in green, orange and red, respectively. The MIF4G domains of UPF2 are in shades of blue, while the partially disordered UPF1-binding domain (U1BD) is denoted as a hatched box. The constructs used in this study are represented by black lines under the respective proteins. B) Analytical SEC and corresponding SDS-PAGE analyses of mixtures of UPF1 and UPF2_S_, in the absence (top panel) and presence of U_15_ RNA (bottom panel). SEC runs of UPF1 and UPF2_S_ in the absence and presence of U_15_ RNA have been included for comparison. Left: Overlay of chromatograms of the mixtures containing the proteins UPF1 and UPF2_S_ (green traces), UPF1 alone (yellow traces) and UPF2_S_ alone (blue traces). In this and all other figures for SEC analysis, solid and dashed lines denote absorbance at 280 nm and 260 nm, respectively. Right: corresponding PAGE analyses of the peak fractions of each run. In addition to SDS-PAGE for visualizing proteins, the bottom panel includes a urea-PAGE analysis of the radiolabeled peak fractions to detect U_15_ RNA. An SDS-PAGE analysis of consecutive fractions of the bottom panel is shown in Supplementary figure 1A. Addition of U_15_ RNA to a mixture of UPF1 and UPF2_S_ does not result in an RNA-bound UPF1-UPF2 complex but rather promotes dissociation of the UPF1-UPF2 complex, resulting in free UPF2_S_, which leads to a broader peak 1, and free UPF1 that binds the U_15_ RNA. C) Analytical SEC analysis of a mixture of a UPF1-UPF2_S_ complex and U_15_ RNA. Top: Overlay of chromatograms of the pre-formed UPF1-UPF2_S_ complex (green traces) and the same complex with U_15_ RNA added (black traces). Bottom: SDS- and urea-PAGE analyses of consecutive SEC fractions in order of increasing retention volume, as indicated. Detection of U_15_ RNA is as described above. Addition of RNA to a stable UPF1-UPF2_S_ complex leads to partial dissociation of the protein complex instead of formation of a stable ternary complex with RNA. D) Native-PAGE analysis of a UPF1-UPF2_S_ complex in the absence and presence of fluorescein-labeled U_15_ RNA, and a UPF1-U_15_ RNA complex in the absence and presence of UPF2_S_. The individual UPF1 and UPF2 proteins as well as the UPF1-UPF2_S_ and UPF1-U_15_ complexes serve as markers for migration of the protein and RNA components on the native gel. Proteins are visualized by staining with Coomassie Brilliant Blue (top panel) and the U_15_ RNA is detected by fluorescence scanning (bottom panel). Asterisks denote the fluorescein label. In solution, UPF1, UPF2_S_ and U_15_ RNA always partition into two binary complexes, UPF1-UPF2_S_ and UPF1-U_15_ RNA. As with analytical SEC (Figure 1C), no ternary UPF1-UPF2_S_-U_15_ complex is detected.

UPF2 is also a multi-domain protein consisting of three MIF4G (middle of eIF4G) domains and a predominantly unstructured C-terminal tail (Figure 1A) (Clerici, Mourao et al. 2009, Clerici, Deniaud et al. 2014). The third MIF4G domain has been shown to bind a number of proteins involved in different aspects of mRNA processing (such as UPF3, Stau1 and the eukaryotic release factor eRF3), while the UPF1-binding region (U1BD in Figure 1A) resides within amino acids 1105 to 1207 (Kadlec, Izaurralde et al. 2004, Clerici, Mourao et al. 2009, Lopez-Perrote, Castano et al. 2016, Gowravaram, Schwarz et al. 2019). The X-ray crystal structure of the UPF1-UPF2 complex shows that the composite U1BD of UPF2 is made up of two distinct secondary structural elements, an α-helix and a β-hairpin, which engage opposite surfaces of the UPF1-CH domain. In-solution NMR studies suggest that the β-hairpin structure is only adopted upon binding to UPF1, indicating that UPF2 also undergoes conformational changes in this process (Clerici, Mourao et al. 2009). Although the available crystal structures of UPF1 bound to RNA and that of a UPF1-UPF2 complex allow insights into activation of UPF1 by UPF2, no experimental structure of a UPF1-UPF2-RNA ternary complex has been determined as yet. As such, our knowledge of assembly of the mRNP that leads to UPF1 activation remains incomplete.

In this study, we investigated the interactions of UPF1, UPF2 and RNA *in vitro*, with an aim to elucidate the mechanism of assembly of the mRNP that leads to UPF1 activation. To our surprise, we found that UPF1 cannot stably associate with UPF2 in the presence of RNA. Addition of RNA dissociates the UPF1-UPF2 complex; conversely addition of UPF2 releases UPF1 from RNA. Previous studies by Chamieh and co-authors also detected a decrease in binding of UPF1 to RNA in the presence of UPF2 and UPF3 (Chamieh, Ballut et al. 2008). Although not obvious from the X-ray crystal structures of RNA-bound UPF1 and the UPF1-UPF2 complex, these observations suggest that the conformation adopted by RNA-bound UPF1 is incompatible with its binding to UPF2, and vice-versa. We present here evidence to suggest that the mutually exclusive interaction *in vitro* is not due to direct competition between UPF2 and UPF1 for RNA but rather indirect effects brought about by conformational changes upon protein-protein/RNA interaction.

## Results and Discussion

The proteins UPF1 and UPF2 have a high binding affinity and readily form a stable complex in solution. We reconstituted a UPF1-UPF2 complex using a construct of UPF1 comprising its CH and helicase domains (referred to hereafter as UPF1, Figure 1A) and a short UPF2 construct encompassing only its MIF4G3 and U1BD domains (UPF2_S_) by mixing together these two proteins in a molar ratio of 1:1.2, with a slight excess of UPF1 to favor complex formation. Analytical size-exclusion chromatography (SEC) of the protein mixture yielded two peaks, a major peak corresponding to the UPF1-UPF2_S_ complex and a smaller peak of the excess UPF1 (Figure 1B, top panel, green traces, and lanes 1 and 2 of the corresponding SDS-PAGE analysis of peak fractions). As a comparison, SEC runs were also performed with UPF1 and UPF2_S_ alone (yellow and blue traces, respectively). To obtain a ternary complex of UPF1, UPF2 and RNA, we added a 15-mer polyuridine RNA (U_15_) to the UPF1-UPF2_S_ mixture and resolved the protein-RNA mixture by analytical SEC (Figure 1B, bottom panel, green traces). We observed a prominent peak for UPF1 bound to RNA, as indicated by the higher absorbance of the peak fractions at 260 nm than 280 nm (Figure 1B, bottom panel, compare peaks and lanes 2 and 4 of SEC and SDS-PAGE analyses) and a broader, asymmetrical peak at approximately the same retention volume as the UPF1-UPF2_S_ complex (Figure 1B, compare peak 1 of top and bottom panels). SDS-PAGE analysis of continuous fractions of this SEC run suggests that the broad peak is an overlap of the peaks of the UPF1-UPF2_S_ complex and UPF2_S_ alone (Supplementary Figure 1A). Notably, the lower absorbance at 260 nm than at 280 nm indicates that RNA is not stably associated with the UPF1-UPF2_S_ complex (although a small amount of U_15_ RNA is detected in lane 1, corresponding to the peak 1 fraction, of the urea-PAGE, Figure 1B, bottom panel). In the presence of U_15_ RNA, not all UPF1 forms a complex with UPF2; some preferentially associates with RNA. SDS-PAGE analyses of the SEC peak fractions of the UPF1/UPF2_S_/RNA mixture and UPF1/UPF2_S_ alone shows that less UPF1-UPF2_S_ complex is formed in the presence of RNA, leaving more UPF1 free to associate with the U_15_ RNA (Figure 1B, compare lane 1 of top and bottom panels).

To ascertain that the inability to isolate a stable UPF1-UPF2-RNA complex was not due to the experimental conditions in which the reconstitution was carried out, we performed analytical SEC with a pre-formed UPF1-UPF2_S_ complex (isolated by preparative SEC) to which U_15_ RNA was added in 1.2-fold molar excess. Instead of a single peak for a UPF1-UPF2_S_-RNA ternary complex, we observed two distinct peaks, one corresponding to UPF1-UPF2_S_ and the other to a UPF1-RNA complex (Figure 1C). A similar observation was made with the UPF1-UPF2_L_ complex that contains a near-full length construct of UPF2 encompassing all three MIF4G domains in addition to the U1BD (Supplementary Figure 1B). We thus infer that addition of U_15_ RNA leads to partial dissociation of the UPF1-UPF2 complex, following which UPF1 binds to RNA.

To corroborate our observations from analytical SEC assays, we performed a native gel analysis using reconstituted complexes of UPF1-UPF2_S_ and UPF1 with a fluorescein-labeled U_15_ RNA (Figure 1D, lanes 6 and 3, respectively). Addition of labeled U_15_ RNA to the UPF1-UPF2_S_ complex led to partial release of UPF2_S_ and formation of a UPF1-U_15_ complex, while addition of UPF2_S_ to the UPF1-RNA complex led to formation of a UPF1-UPF2_S_ complex, due to the availability of UPF1 after partial dissociation of RNA from the UPF1-U_15_ RNA complex (Figure 1D, lanes 7 and 8 respectively). Similarly, an electrophoretic mobility shift assay (EMSA) of increasing concentrations of the UPF1-UPF2_L_ complex with a radiolabeled U_15_ RNA shows appearance of a predominant shifted band corresponding to a UPF1-U_15_ complex and a weaker band corresponding to a UPF2_L_-U_15_ complex. No super-shifted band corresponding to a ternary UPF1-UPF2_L_-U_15_ complex was obtained (Supplementary Figure 1C). Taken together, our observations clearly show that a stable complex of UPF1, UPF2 and RNA cannot be reconstituted *in vitro* as binding of UPF1 to RNA is incompatible with its interaction with UPF2.

In cells, UPF1 is approximately 10-fold more abundant than UPF2 and can bind along the length of an mRNA transcript (Hein, Hubner et al. 2015, Cho, Cheveralls et al. 2022). In the context of NMD, UPF2 is associated with the EJC via UPF3. A cryo-EM structure of the EJC-UPF3-UPF2-UPF1 complex shows UPF1 positioned towards the 3’-end of the RNA (Melero, Buchwald et al. 2012). However, UPF1 was also shown to be involved in NMD target selection, using its ATPase activity for discriminating between target and non-target mRNAs (Lee, Pratt et al. 2015). This step presumably occurs directly at or in close proximity of the premature termination codon (PTC), upstream of the EJC, where the translating ribosome is stalled. It is possible that binding of UPF2 temporarily displaces UPF1 from RNA and repositions it downstream of the EJC to remodel the 3’-end of the mRNP. To gain deeper insight into displacement of UPF1 from RNA by UPF2, we performed fluorescence anisotropy assays, where increasing amounts of UPF2_S_ were titrated into a mixture of UPF1 and U_12_ RNA that was labeled with 6-FAM at its 5’-end. In the absence of UPF2_S_, the UPF1-RNA mixture showed high fluorescence anisotropy, as expected. Increasing UPF2_S_ concentrations led to a decrease in fluorescence anisotropy, which can be attributed to release of UPF1 from the labeled RNA (Figure 2A). Interestingly, the fluorescence anisotropy at the highest concentrations of UPF2_S_ tested does not go down to 0 (the value corresponding to free RNA in solution) but plateaus off at a relative value of 0.23, suggesting that some UPF1 remains bound to the RNA. This is in accordance with our observations from native PAGE analysis (Figure 1D), where addition of UPF2_S_ does not result in complete dissociation of the UPF1-RNA complex.

**Figure 2.**
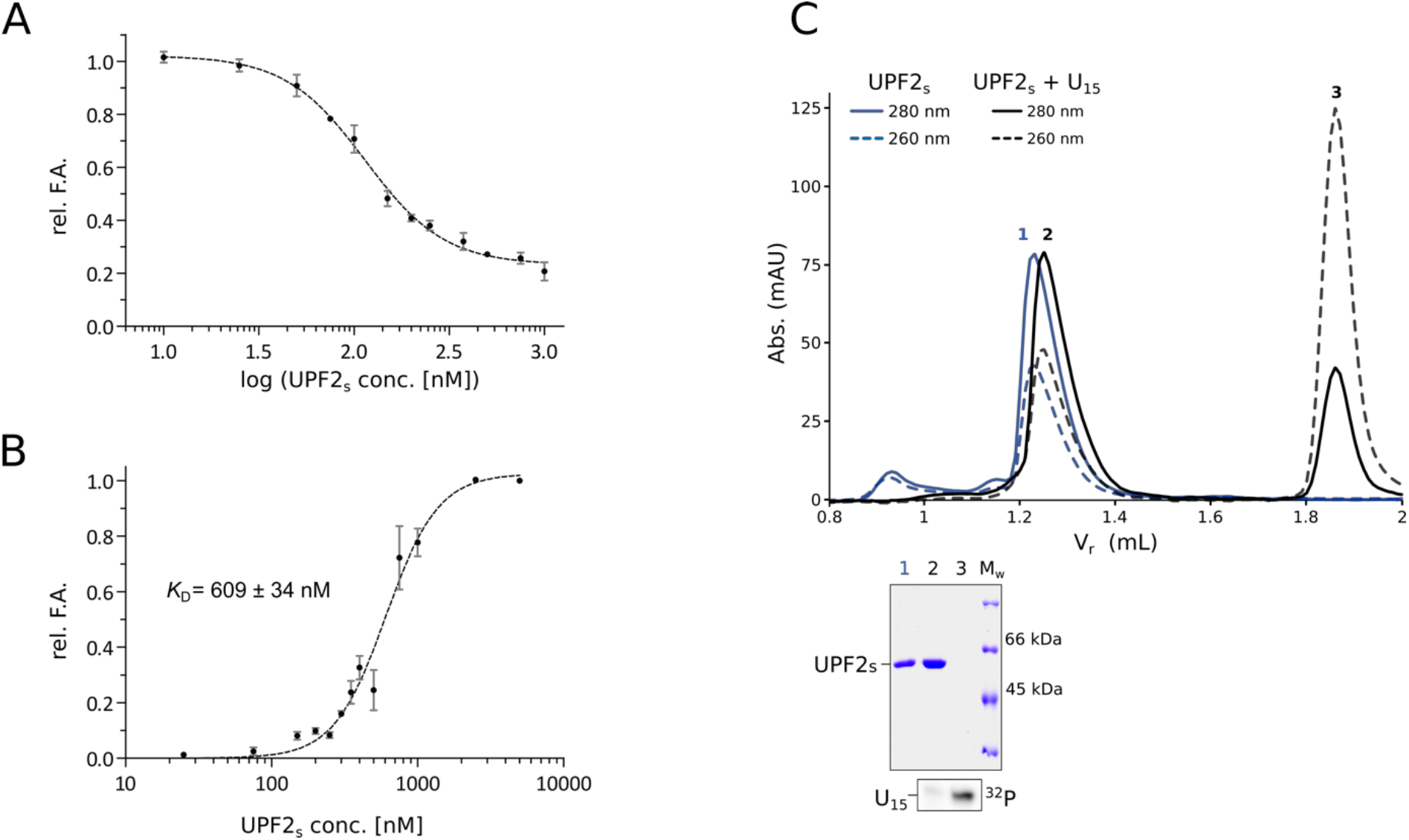
RNA binding properties of UPF2_S_ and its impact on binding of UPF1 to RNA. A) Fluorescence anisotropy competition assay to determine how addition of UPF2_S_ affects binding of UPF1 to 6-FAM-U_12_ RNA. Titration of increasing amounts of UPF2_S_ into a constant amount of UPF1-U_12_ RNA mixture results in partial displacement of UPF1 from RNA. The data points and errors bars of this and all other fluorescence anisotropy experiments are the mean and standard deviation of at least 2 independent experiments. B) Fluorescence anisotropy assay to determine the dissociation constant (*K*_D_) of the UPF2_S_-U_12_ RNA interaction. The error associated with the *K*_D_ is its standard deviation (SD). UPF2_S_ binds U_12_ RNA with a modest affinity, which is an order of magnitude lower than the RNA-binding affinity of UPF1. C) Analytical SEC analysis of binding of UPF2_S_ to U_15_ RNA. Top and bottom panels show the overlay of chromatograms of UPF2_S_ without (blue traces) and with RNA (black traces), and the corresponding PAGE analyses of peak fractions of each run, respectively. UPF2_S_ does not form a stable complex with RNA.

We next tested if the release of UPF1 from RNA in the presence of UPF2 is a consequence of direct competition for RNA binding. Although previous studies demonstrated binding of UPF2 to RNA, the affinity of this interaction remains unknown (Kadlec, Izaurralde et al. 2004). Therefore, we carried out fluorescence anisotropy assays of UPF2_S_ with 6-FAM-U_12_ RNA (Figure 2B) and determined a dissociation constant (*K*_D_) of approximately 600 nM, an order of magnitude higher than the *K*_D_ of apo-UPF1 for RNA (∼ 50 nM). Furthermore, an analytical SEC assay of UPF2_S_ with the U_15_ RNA showed that it does not form a stable complex with RNA (Figure 2C), which is consistent with the weak binding of UPF2_S_ to RNA observed in EMSA (Supplementary Figure 1C). These observations suggest that it is unlikely that the UPF2-mediated dissociation of UPF1 from RNA is a result of preferential binding of UPF2_S_ to RNA, but rather due to UPF2-induced conformational rearrangements within UPF1, which are incompatible with its RNA binding. This also explains the observation that addition of RNA leads to disruption of a UPF1-UPF2 complex and that no ternary complex of UPF1, UPF2 and RNA can be isolated *in vitro* (Figure 1C).

To test this hypothesis, we used a short construct of UPF2 (UPF2-U1BD, Figure 1A) that spans the UPF1 binding region but lacks the MIF4G3 domain that was earlier shown to bind RNA. The affinity of UPF2-U1BD for RNA is far lower than that of UPF2_S_, precluding determination of a *K*_D_ for RNA binding by fluorescence anisotropy (Figure 3A). Nevertheless, addition of increasing amounts of UPF2-U1BD to a mixture of UPF1 and 6-FAM-U_12_ RNA led to a steady decrease in fluorescence anisotropy, corresponding to a release of UPF1 from RNA (Figure 3B). Conversely, addition of U_15_ RNA to a complex of UPF1 with UPF2-U1BD led to partial dissociation of the complex in analytical SEC, as indicated by the appearance of a peak corresponding to UPF1-U_15_ RNA (Supplementary Figure 1D). It appears that UPF2-U1BD, without interacting with RNA, mediates the same effect as UPF2_S_ on RNA binding of UPF1, substantiating the argument that the inability of UPF1 to concomitantly interact with UPF2 and RNA in a stable manner is not due to competition between the two proteins for RNA. To corroborate this observation, we tested the ability of a UPF2 construct lacking the C-terminal MIF4G3 and UPF1-binding domains (UPF2-MIF4G1-2) to bind RNA and to displace UPF1 from RNA. We find that UPF2-MIF4G1-2 binds U_12_ RNA with an affinity comparable to that of UPF2_S_ but unlike UPF2_S_, it does not displace UPF1 from RNA (Figures 3A and 3B). We conclude that dissociation of UPF1 from RNA upon addition of UPF2, and vice-versa, stems from rearrangements in protein-protein/RNA interactions upon addition of the second binding partner and prevents formation of a stable ternary UPF1-UPF2-RNA complex.

**Figure 3.**
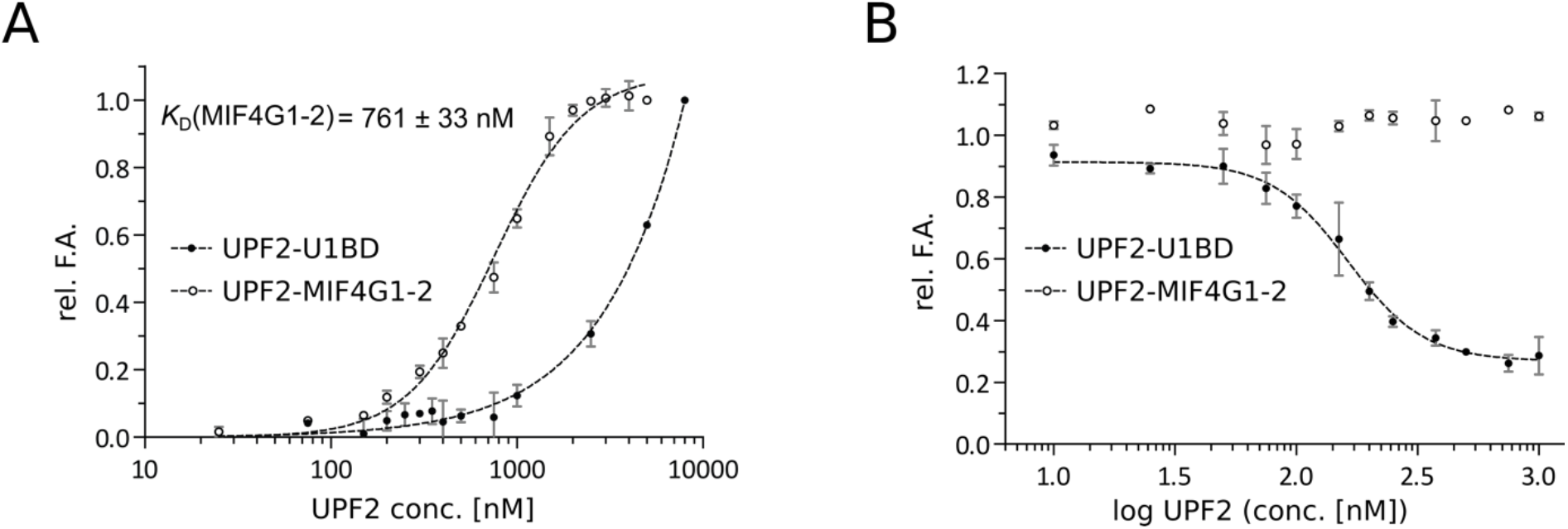
Displacement of UPF1 from RNA by UPF2 is not due to competition in RNA-binding. A) Quantitative measurements of RNA-binding affinities of UPF2 constructs comprising (UPF2-U1BD) and lacking the UPF1-binding site (UPF2-MIF4G1-2) by fluorescence anisotropy using 6-FAM-labeled U_12_ RNA. The *K*_D_ of UPF2-MIF4G1-2 is reported along with its SD. The affinity of UPF2-MIF4G1-2 for RNA is comparable to that of UPF2_S_ (open circles) while UPF2-U1BD does not show any appreciable affinity for RNA (filled circles). B) Fluorescence anisotropy competition assays to determine the effect of UPF2-U1BD and UPF2-MIF4G1-2 on the UPF1-RNA interaction. The UPF2-U1BD protein that binds UPF1, but not RNA, is capable of displacing UPF1 from RNA (filled circles), whereas the MIF4G1-2 construct that binds RNA but lacks the UPF1-binding motif has no impact on RNA-binding by UPF1 (open circles).

## Conclusions

The NMD pathway involves the assembly and disassembly of several protein/RNA complexes to discern mRNA transcripts as *bona fide* substrates and subsequently remodel mRNPs and completely degrade target mRNAs (reviewed in (Karousis and Muhlemann 2019, Kishor, Fritz et al. 2019, Kurosaki, Popp et al. 2019)). As a core component of the NMD pathway, UPF1 is involved in several of these complexes, often through transient interactions (Lavysh and Neu-Yilik 2020). Examples of UPF1-centric complexes in NMD are the SURF (SMG1-UPF1-eRFs) complex which is assembled on the PTC-stalled ribosome and the decay-inducing (DECID) complex that is formed upon association of SURF with UPF2-3-bound EJC (Yamashita, Ohnishi et al. 2001, Kashima, Yamashita et al. 2006). Additionally, both phosphorylated and unphosphorylated UPF1 interact with the endonuclease SMG6 and phosphorylated UPF1 also engages the SMG5/SMG7 heterodimer, which in turn bridges the target mRNP to the deadenylation machinery (Okada-Katsuhata, Yamashita et al. 2012, Loh, Jonas et al. 2013, Chakrabarti, Bonneau et al. 2014, Nicholson, Josi et al. 2014). The involvement of SMG6 in NMD depends both on UPF1 as well as binding of SMG5/SMG7 to phospho-UPF1, indicating a complex interplay of interactions at work in this pathway (Boehm, Kueckelmann et al. 2021). It therefore appears intuitive that many of these interactions must be short-lived in order to facilitate progression of NMD and efficient degradation of the nonsense mRNA.

In this study, we show that UPF1 engages UPF2 and RNA in mutually exclusive interactions. We investigate this in detail and provide an underlying mechanism for our observations and that of Chamieh and co-workers, where binding of UPF1 to RNA was weakened in the presence of UPF2. These observations are somewhat counter-intuitive to our understanding of stimulation of UPF1’s helicase and RNA-dependent ATPase activities by UPF2. Activation of UPF1 by UPF2 is a crucial step in NMD as the helicase has a low basal catalytic activity *in vitro* in the absence of its binding partners. It is widely speculated that activation of UPF1 occurs in the context of the EJC-UPF3-2-1 complex. The cryo-EM structure of the EJC-UPF complex shows UPF1 positioned towards the 3’-end of the EJC-bound RNA, poised to translocate in the 5’-3’ direction (Melero, Buchwald et al. 2012). However, it is not clear from structural and biochemical studies whether activation of UPF1 occurs early on in the NMD pathway (immediately after association with EJC-UPF3-2) or at a later stage, just prior to degradation. It is possible that association of UPF1 with the EJC-UPF3-2 complex leads to its activation as well as dissociation from the mRNA, while still being tethered to the mRNP via protein-protein interactions. Once activated UPF1 rebinds the mRNA, it would dissociate from the EJC-UPF3-2 complex and translocate along the mRNA towards its 3’-end. Alternatively, EJC-independent UPF2 could bind UPF1, activate it and dissociate from UPF1 rapidly, leaving it free to rebind the mRNA. This raises the question of how UPF2 is recruited to the nonsense-mRNP independent of the EJC. We present evidence for binding of UPF2 to RNA with a modest affinity, although this interaction is not as stable as that of UPF1 with RNA. It is not known if UPF2 associates with other NMD components, apart from UPF1 and UPF3.

Recent studies on another UPF1-mediated decay pathway, Staufen-mediated mRNA decay (SMD) showed the involvement of UPF2, where it acts as an adaptor between UPF1 and the double-stranded (ds) RNA-binding protein Stau1 and activates UPF1 within this complex (Gowravaram, Schwarz et al. 2019). Although the ability of the UPF1-UPF2-Stau1 complex to interact with dsRNA has been shown in this study, it would be worthwhile to investigate if the protein complex can stably associate with dsRNA and if UPF2 also affects the RNA-binding properties of Stau1. How UPF1 activation by UPF2 is achieved in the various decay pathways while keeping the helicase associated with the target mRNP therefore remains a conundrum that requires further investigation.

### Experimental Procedures

#### Protein expression and purification

All human UPF1 and UPF2 constructs used in this study were expressed as 6x-His or His-Thioredoxin (Trx) fusions in *Escherichia coli* BL21 (*DE3*) STAR pRARE cells at 18 °C for at least 15 hours. Cells expressing recombinant proteins were lysed using lysis buffer (50 mM Tris-HCl pH 7.5, 500 mM NaCl, 10 % glycerol, 1 mM MgCl_2_, 1 µM ZnCl_2_, 0.1 M urea and 10 mM imidazole,), supplemented with protease inhibitors (1 mM PMSF) and DNase I. The proteins were isolated from the crude lysate by Ni^2+^-affinity chromatography and washed successively with lysis buffer, chaperone wash buffer (50 mM Tris-HCl pH 7.5, 1 M NaCl, 10 % glycerol, 10 mM MgCl_2_, 50 mM KCl, 1 µM ZnCl_2_, 2 mM ATP and 10 mM imidazole) and low salt wash buffer (50 mM Tris-HCl pH 7.5, 150 mM NaCl, 10 % glycerol, 1 mM MgCl_2_, 1 µM ZnCl_2_, and 10 mM imidazole) gradually. Finally, target proteins were eluted with nickel column elution buffer (50 mM Tris-HCl pH 7.5, 150 mM NaCl, 10 % glycerol, 1 mM MgCl_2_, 1 µM ZnCl_2_, and 300 mM imidazole). The affinity tags on UPF1 and UPF2 constructs were not removed. UPF1, UPF2_L_ and UPF2_S_ were subjected to a further purification step using a HiTrap Heparin Sepharose HP column (GE Healthcare) and heparin buffers A (20 mM Tris-HCl pH 7.5, 10 % glycerol, 1 mM MgCl_2_, 1 µM ZnCl_2_, and 2 mM DTT) and B (20 mM Tris-HCl pH 7.5, 1 M NaCl, 10 % glycerol, 1 mM MgCl_2_, 1 µM ZnCl_2_, and 2 mM DTT). Proteins were eluted from the column using a linear concentration gradient of NaCl. All proteins including UPF2-U1BD were purified by a final size exclusion chromatography step (using Superdex 75 or Superdex 200 columns, GE Healthcare) in SEC buffer (20 mM Tris-HCl pH 7.5, 150 mM NaCl, 5 % glycerol, 1 mM MgCl_2_, 1 µM ZnCl_2_, and 2 mM DTT).

Complexes of UPF1 and various UPF2 constructs were formed by mixing together the two proteins in a 1:1.2 molar ratio (with UPF1 in excess) overnight at 4 °C, followed by SEC on a Superdex 200 column.

#### Analytical SEC

700 pmol of the single proteins (UPF1 or UPF2 alone), UPF1-UPF2 complexes or the mixtures of approximately equimolar amounts of UPF1 and UPF2 were mixed with 700 pmol of a 15-mer poly(U)-RNA (U_15_) (Eurofins Genomics) to a final volume of 50 µL in A-SEC buffer (20 mM HEPES pH 7.5, 100 mM NaCl, 5 % glycerol, 1 mM MgCl_2_, 1 µM ZnCl_2_, 2 mM DTT) and incubated on ice overnight. Individual proteins and protein-RNA complexes were resolved on a Superdex 200 Increase 3.2/300 column (GE Healthcare). The peak fractions were analyzed by SDS-PAGE, followed by staining with Coomassie Brilliant Blue. Peak fractions for analytical SEC runs performed with U_15_ RNA were radiolabeled with [γ-^32^P]-ATP, as described below and visualized on 15% urea-PAGE by autoradiography. The curves of chromatograms in the same figure were normalized manually for intuitive comparison.

### Sample labelling and electrophoretic mobility shift assays

2 µl of peak fractions from indicated SEC runs were treated with [γ-^32^P]-ATP (Hartmann Analytic GmbH) and T4 polynucleotide kinase (Molox GmbH) in PNK A buffer (Thermofisher) for 1.5 h at 37 °C in order to 5’-end label the U_15_ RNA therein. Excess [γ-^32^P]-ATP was separated from the labelled RNA by purification on a G25 spin column (GE HealthCare).

For electrophoretic mobility shift assays (EMSA) 10 pmol U_15_ RNA (Eurofins Genomics) was radiolabelled at the 5’-end with [γ-^32^P]-ATP and subsequently purified as described above. 0.2 pmol of radiolabelled RNA was incubated with 150 nM UPF1, 150 nM UPF2 or a pre-formed UPF1-UPF2 in assembly buffer (20 mM HEPES, 100 mM NaCl, 0.1 % NP-40, 0.5 % Glycerol, 1 mM DTT, 0.5 mM EDTA) in a total volume of 10 µl for 30 minutes on ice. Samples were resolved on a 5 % native PAGE that was run at 160 V for 2.5 hours at 4 °C. Bands containing RNA were visualized via autoradiography using a phosphor-imager (GE Healthcare).

#### Native gel analysis

A UPF1-U_15_ RNA complex was formed on a preparative scale by mixing 100 µg UPF1 protein with a 1.2-fold molar excess of fluorescein-labelled U_15_ RNA, followed by size-exclusion chromatography on a 2.4 mL Superdex 200 column. The peak fraction containing both UPF1 and U_15_ RNA was used for the native gel analysis. 2 µg of individual proteins and 4 µg of complexes were used in each case. Proteins were mixed with a 1.2-fold molar excess of fluorescein-labelled U_15_ RNA and incubated at room temperature for 30 minutes. Thereafter, samples were reconstituted in native gel sample buffer and directly analysed on a 4-20% Tris-Glycine native gel (ThermoFisher Scientific). Proteins were visualised by staining with Coomassie Brilliant Blue and the U_15_ RNA was detected by fluorescence scanning using a Typhoon scanner (GE Healthcare).

#### Fluorescence anisotropy

To determine the affinity of UPF2 constructs for RNA, 10 nM of a 12mer poly(U)-RNA (U_12_) labelled with 6-FAM at its 5’-end was mixed with increasing concentrations of human UPF2_S_, UPF2-MIF4G1-2 and UPF2-U1BD in FA-binding buffer (20 mM HEPES pH 7.5, 100 mM NaCl, 1 mM MgCl_2_, 100 μg/mL BSA). 40 μL of each sample were transferred to a black 384-well plate (PerkinElmer OptiPlate 384-F) and fluorescence polarization was measured with a Tecan Spark plate reader at 25°C. The reading obtained in the absence of UPF2 (RNA alone sample) was considered as background and subtracted from all fluorescence polarization (FP) values. Fluorescence anisotropy was calculated from fluorescence polarization using the formula 2·FP/(3-FP) and normalized against the value obtained for the highest UPF2 concentration. The data shown are an average of at least 2 independent experiments and were fitted to an equation representing one site - specific binding with Hill slope in GraphPad Prism 5.00. Error bars and the error associated with the reported *K*_D_ denote standard deviation (SD).

In order to determine the effect of different UPF2 constructs on RNA-binding of UPF1, 70 nM hUPF1 and 10 nM of 6-FAM-labelled U_12_ RNA were mixed with increasing concentrations of human UPF2_S_, UPF2-MIF4G1-2 and UPF2-U1BD in FA-binding buffer. Fluorescence polarisation was recorded and fluorescence anisotropy was calculated as described above. The data, an average of 3 independent experiments (except for UPF2-MIF4G1-2, where only 2 independent experiments were performed), were fitted, when possible, to an equation describing dose-dependent inhibition (log(inhibitor) vs. response - variable slope) in GraphPad Prism 5.00. As above, data point and error bars represent the mean and standard deviation of at 3 independent experiments.

## Data availability

All data for this manuscript are contained within the main article and supplementary information.

## Acknowledgements

We thank Elena Conti for support at the initial stage of this project, Markus Wahl for the gift of 6-FAM-labeled U_12_ RNA and Juliane Metzke for purification of the UPF2-MIF4G1-2 protein.

## Author Contributions

S.C. – conceptualization, funding acquisition, supervision; G.X. – protein purification and SEC analysis; V.M. and M.P. – fluorescence anisotropy; A.A. and M.P. – EMSAs; U.J. – native PAGE analysis; A.A. and S.C. – writing - original draft; All authors – writing - review and editing.

