## Supplementary Figure for "Modulation of RNA binding properties of the RNA helicase UPF1 by its activator UPF2"

### Supplementary Information

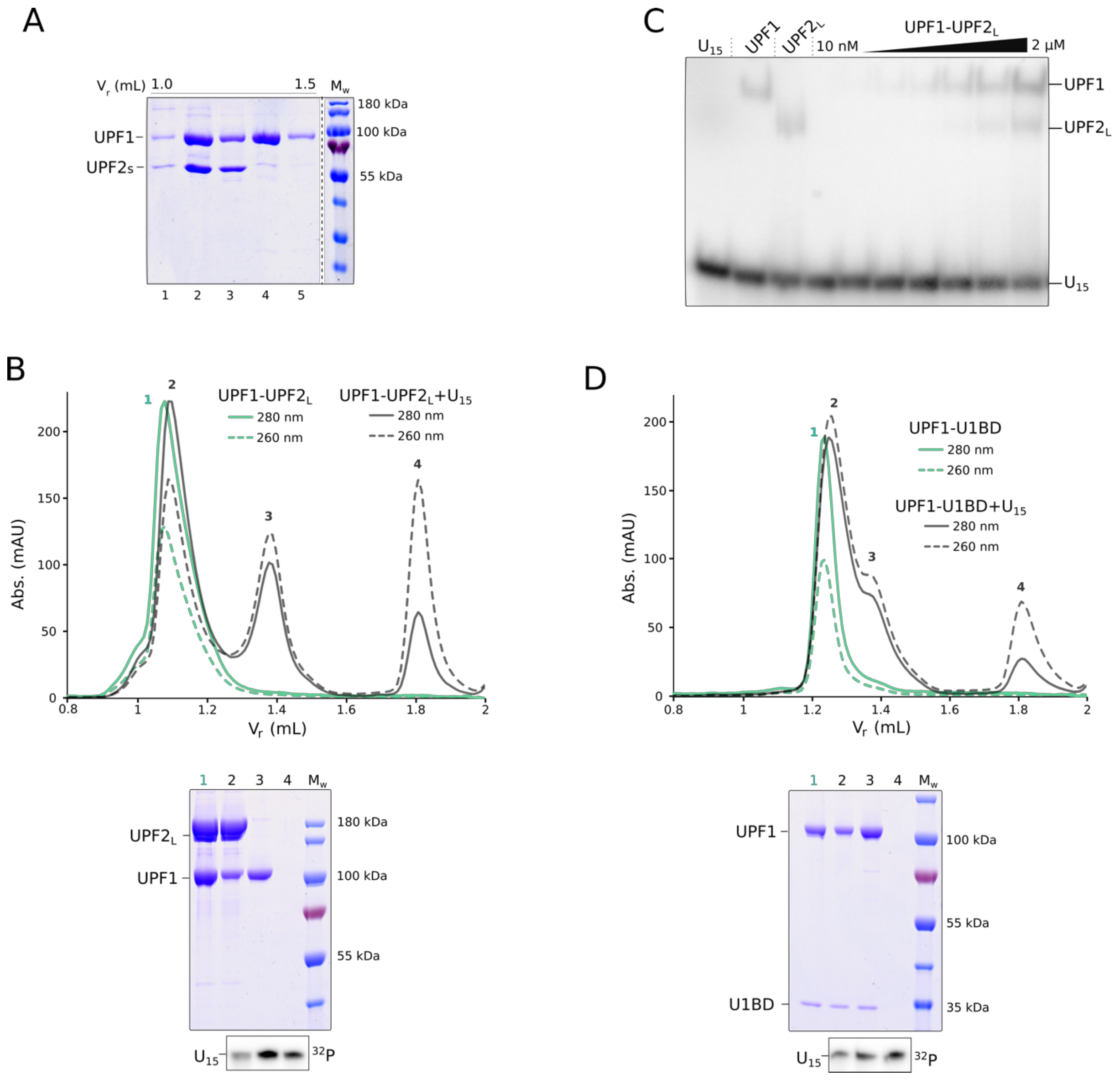


**Supplementary Figure 1: Impact of different UPF2 constructs on RNA-binding by UPF1**

A) Coomassie-stained SDS-PAGE analysis of the peak fractions of the UPF1-UPF2$_S$ complex + $U_{15}$ RNA SEC run, shown in Figure 1B. Fractions are loaded in ascending order of retention volume. A careful comparison of lanes 2 and 3 reveals a change in the relative intensity of the UPF1 and UPF2$_S$ bands, which indicate partial dissociation of the UPF1-UPF2$_S$ complex. Dissociation of the UPF1-UPF2$_S$ complex releases free UPF2$_S$ which co-migrates with the complex peak (Peak 1 in Figure 1B, bottom panel) and UPF1 that binds the $U_{15}$ RNA (Peak 2). C) Electrophoretic mobility shift assay of the UPF1-UPF2$_S$ complex with $^{32}$P-labeled $U_{15}$ RNA. The absence of a super-shifted band in comparison to RNA-bound UPF1 and UPF2 suggests that no ternary complex of UPF1-UPF2$_S$-$U_{15}$ is formed. As seen in SEC analysis, addition of RNA to the UPF1-UPF2$_S$ complex causes it to dissociate. B) and D) Analytical SEC and corresponding PAGE analyses of UPF1-UPF2$_L$ and UPF1-UPF2-U1BD complexes in the presence of RNA. Proteins are visualized by staining with Coomassie Brilliant Blue while $U_{15}$ RNA is detected by radiolabeling the peak fractions with $^{32}P$. Addition of RNA to both complexes leads to their dissociation, as indicated by appearance of a UPF1-RNA peak (peak 3 in both chromatograms). As in figures in the main text, solid and dashed lines denote absorbance at 280 nm and 260 nm, respectively. The higher absorbance at 260 nm and appearance of $U_{15}$ RNA in peak 2 is not due to RNA binding of the UPF1-UPF2-U1BD complex but due to partial overlap of the complex peak with that of RNA-bound UPF1 (peak 3).
